# Using big sequencing data to identify chronic SARS-Coronavirus-2 infections

**DOI:** 10.1101/2023.07.16.549184

**Authors:** Sheri Harari, Danielle Miller, Shay Fleishon, David Burstein, Adi Stern

**Author notes:** Equal contribution (Sheri Harari, Danielle Miller).

## Abstract

The evolution of SARS-Coronavirus-2 (SARS-CoV-2) has been characterized by the periodic emergence of highly divergent variants, many of which may have arisen during chronic infections of immunocompromised individuals. Here, we harness a global phylogeny of ∼11.7 million SARS-CoV-2 genomes and search for clades composed of sequences with identical metadata (location, age, and sex) spanning more than 21 days. We postulate that such clades represent repeated sampling from the same chronically infected individual. A set of 271 such chronic-like clades was inferred, and displayed signatures of an elevated rate of adaptive evolution, in line with validated chronic infections. More than 70% of adaptive mutations present in currently circulating variants are found in BA.1 chronic-like clades that predate the circulating variants by months, demonstrating the predictive nature of such clades. We find that in chronic-like clades the probability of observing adaptive mutations is approximately 10-20 higher than that in global transmission chains. We next employ language models to find mutations most predictive of chronic infections and use them to infer hundreds of additional chronic-like clades in the absence of metadata and phylogenetic information. Our proposed approach presents an innovative method for mining extensive sequencing data and providing valuable insights into future evolutionary patterns.

## Introduction

The evolution of SARS-Coronavirus-2 (SARS-COV-2) has been punctuated by the periodic emergence of variants that are highly genetically divergent compared to the circulating variants at the time of their emergence. Some of these variants were found to be more transmissible than their predecessor variants, leading to patterns of rapid displacement of one variant by another. Specific variants of concern (VOCs) were designated by the World Health Organization (WHO) when there was concrete evidence that a variant posed an increased risk to public health. The nomenclature of SARS-COV-2 variants has been challenging ^1^; here we refer to VOCs mainly based on the Greek letters assigned by PANGO ^2^ and WHO ^1^. When more precise nomenclature is necessary, we rely on the classification of Nextstrain clades ^3^. As of the end of 2020, a series of VOCs created patterns of global displacement, namely Alpha, Delta, Omicron BA.1, Omicron BA.2, and lastly Omicron BA.5. More recently, milder global changes have been occurring, as at the time of writing there are multiple variants co-circulating globally, and these variants are usually not dramatically divergent compared to their predecessors.

One of the leading hypotheses regarding the origin of highly divergent SARS-COV-2 variants is that they emerged in chronically infected individuals ^4^. Chronic infections (also called persistent or prolonged infections) are herein defined as infections where there is evidence of actively replicating virus for more than 21 days, and often such infections may last months or even more than a year ^5^. To date, all chronic infections were found to occur in immunocompromised individuals suffering from one of four categories: hematologic cancer, AIDS, transplant patients or autoimmune patients^6–10^. Chronic infections should not be confused with long COVID where symptoms persist but not necessarily active viral replication.

The hypothesis on the origin of VOCs is supported by the fact that different combinations of VOC lineage-defining mutations, particularly in the spike gene, are observed across chronic infections ^5^. It is thought that the partially functioning immune system in chronically infected individuals, coupled with treatments, pose strong selection pressure that leads to rapid adaptation of the virus. This contrasts with a slower rate of adaptive evolution observed along transmission chains of acute infections. Specifically, the very narrow transmission bottleneck found to occur during transmission in SARS-COV-2 is expected to dramatically dampen the rate of adaptive evolution during community transmission ^11–13^.

Despite the increased understanding that chronic SARS-COV-2 infections may significantly contribute to the evolution of the virus, our understanding of the dynamics of these infections and how they impact global evolutionary patterns is limited and mostly contingent on isolated case reports. Our and other previous meta-analyses ^5, 14^ were mostly limited to the pre-VOC era and were based on a small number of cases (but see ^15^). We posit here that the ever-growing database of millions of SARS-COV-2 genomes likely harbors many sequences derived from chronic infections. We suggest an approach to mine these data by harnessing the phylogeny and associated metadata submitted with the sequencing data (age, sex, location, and dates).

Notably, sequences are usually not explicitly identified as originating from chronically infected individuals or from the same person. We reasoned that most often, sequences derived from chronically infected individuals will display as monophyletic clades, i.e., all sequences derived from the ancestral node of the clade are from the same individual^7, 9, 10^. This reflects the following assumptions: (i) sometimes, chronically infected individuals will be serially sampled and sequenced, (ii) sequences derived from the same individual will be very similar, and (iii) most chronically infected patients do not create onward transmission chains ^5, 15^ (otherwise, the clade would not be monophyletic). We accordingly mined a global phylogeny of over 11 million sequences and inferred 271 inferred chronic-like monophyletic clades and a set of control clades derived from transmission chains. By comparing chronic-like clades to controls, we were able to obtain insights regarding the different evolutionary dynamics of chronic versus acute infections.

Analysis of data at the scale described herein poses many challenges, including missing data and computational limitations. Recently, language models have gained widespread popularity due to their ability to tackle such large volumes of data and have impacted research in diverse areas of biological sciences, including virology ^16–18^. To this end, we developed language models to effectively capture the distinct mutational patterns exhibited by chronic-like clades in comparison to control clades. This enabled us to identify specific mutations that exhibit a stronger association with chronic-like clades. Finally, we used this model to infer chronic infections, in the absence of metadata and in the absence of information on the phylogenetic structure of the sequences. Our analysis allowed us to demonstrate that in-depth mining of the huge volume of SARS-COV-2 sequences is highly informative and that chronic-like clades are predictive of future evolution.

## Results

We began by analyzing over 11.7 million sequences of SARS-COV-2 and their associated phylogeny after stringent quality control filtering (Methods). The phylogeny was mined for monophyletic clades where all sequences share the same metadata (defined here as location, age, and sex) and where the sampling dates of the sequences span more than 21 days, resulting in 271 chronic-like clades (also denoted as cases). In addition, we constructed a set of positive and negative control clades. The set of positive clades was composed of thirty-two *bona fide* chronic infections derived from published case reports for which sequencing, metadata and clinical information were available (Methods). The set of negative control clades was generated by sampling 15,163 monophyletic clades that did not share the same metadata (Methods) (Fig. S1), to ensure that these clades most likely represent transmission chains from acute infections. In the analyses below, when comparing between cases and controls, we performed stratified sampling from this set of controls to maintain identical sample sizes of *n*=271, and to maintain similar distributions of clade sizes and background variants between cases and controls (Methods).

The 271 chronic-like clades were distributed across all the major variants detected till September 2022 (Fig. 1a, Table S1) and originated from a wide range of countries, with a small bias towards Europe (Table S2). We first set out to test whether we could find evolutionary signatures that would suggest that the set of chronic-like clades found herein are enriched for valid chronic infections.

**Figure 1.**
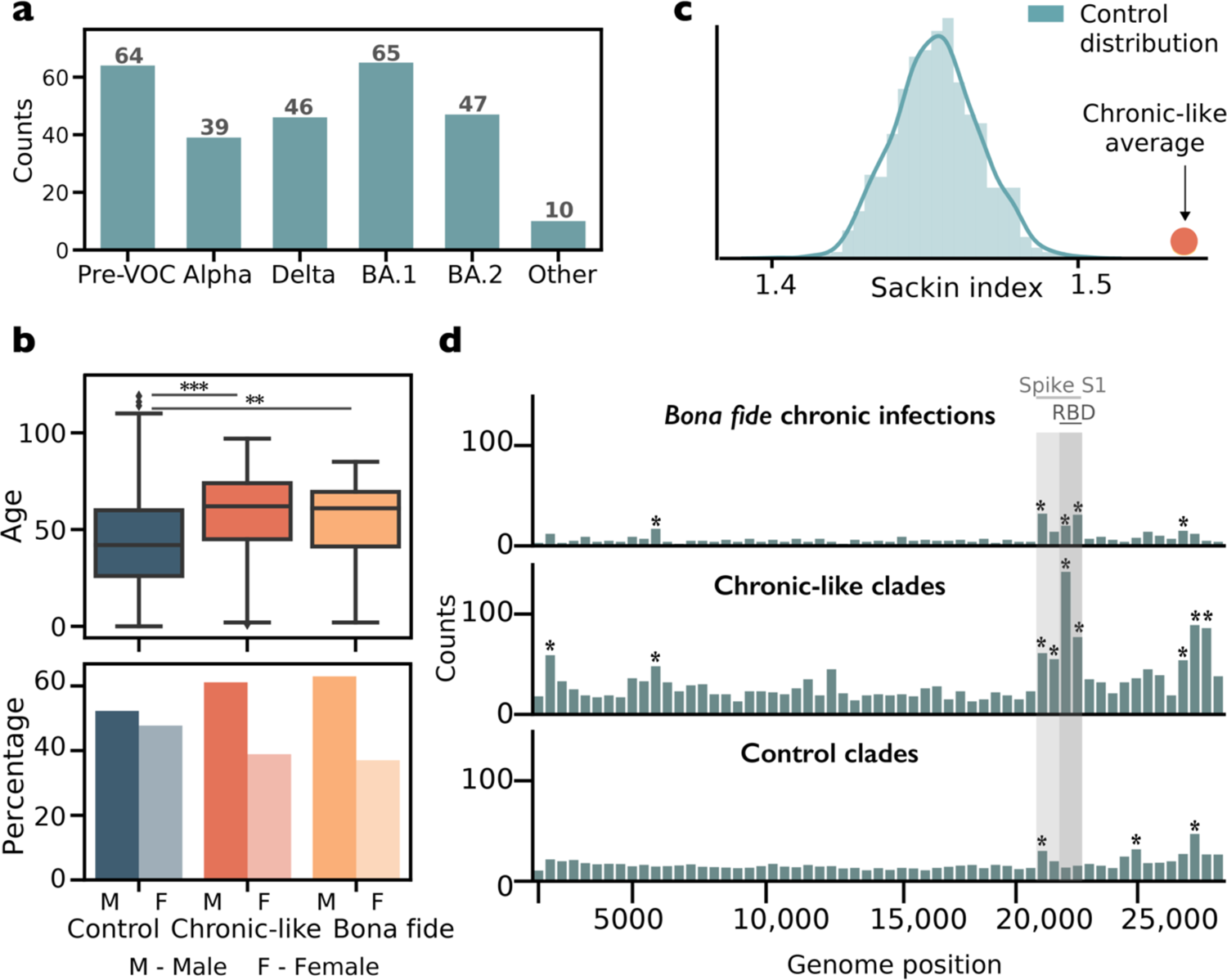
Characteristics of chronic-like clades compared to control clades and *bona fide* chronic infections. (a) The number of chronic-like clades stratified by background variant. Pre-VOC refers to any variant that was dominant before the emergence of Alpha. (b) The distribution of Sackin index values over repeated stratified sampling of n=271 control clades, with the orange circle representing the average Sackin index of the chronic-like clades. (c) Distribution of age and percentage of male/female shown for control clades, chronic-like clades and *bona fide* chronic infections. (d) Distributions of substitutions along the SARS-CoV-2 genome observed across all *bona fide* chronic infections, chronic-like clades, and control clades. Substitutions are counted in bins of 500 nucleotides. Asterisks mark bins significantly enriched for more substitutions using a one-tailed binominal test, after correction for multiple testing (*p* < 0.0001).

We first compared basic demographic features of cases and controls, namely age and sex. We found that on average chronic-like clades were characterized by older age, and a higher proportion of males as compared to the control clades (Fig. 1b). This trend highly resembles the *bona fide* chronic infections. This is in line with a higher tendency for older males to suffer from hematologic cancers, which represented the majority of chronic infections in our previous study^5^. Additionally, these individuals are more likely to present with COVID-19 deteriorations or hospitalizations ^19, 20^, possibly leading to a higher chance of being sequentially sampled.

Next, we compared the distribution of substitutions found in the sets of *bona fide* chronic infections, chronic-like clades, and control clades, in bins of 500 bases along the genome (Fig. 1d). Our results show that both *bona fide* chronic infections and chronic-like clades are enriched for spike mutations, particularly in the S1 subunit of the spike protein. This is in line with repeated selection observed at this region ^21^, most likely for antibody evasion and/or enhanced ACE2 binding. In contrast, the distribution of substitutions along the control clades is most often uniform, with the exception of some enrichment in genes in the 3’ region of the genome, found across all sets, and a small enrichment in the spike N-terminal domain. Indeed, it has been shown that many genes in the 3’ region of the genome are under relaxed selection and tend to accumulate more mutations ^22^.

We went on to analyze the tree topology of the chronic-like clades. When comparing inter-host to intra-host evolution, we expect more ladder-like trees in the latter, reflecting adaptive evolution and stepwise accumulation of beneficial mutations over time ^23^. On the other hand, acute infection transmission chains are expected to be characterized by superspreading events that lead to a star-like phylogeny. These two features are captured by the Sackin index of a tree, with higher values for ladder-like trees and lower values for star-like trees. In line with this assumption, we found that on average our chronic-like clades bore significantly higher Sackin index values as compared to controls (*p <* 10^-4^, permutation test, Fig. 1c).

We present three examples of chronic-like clades (Fig. 2a-c) as well as an illustration of a control clade (Fig. 2d). As we describe herein, many chronic-like clades exhibit rapid evolution, especially in the spike gene. However, it is important to note that we also observed that in some chronic-like clades there was not necessarily dramatic evolution (e.g., Fig. 2c). This is in line with our previous report, where we observed that chronic infections varied in their inferred rate of adaptive evolution ^5^.

**Figure 2.**
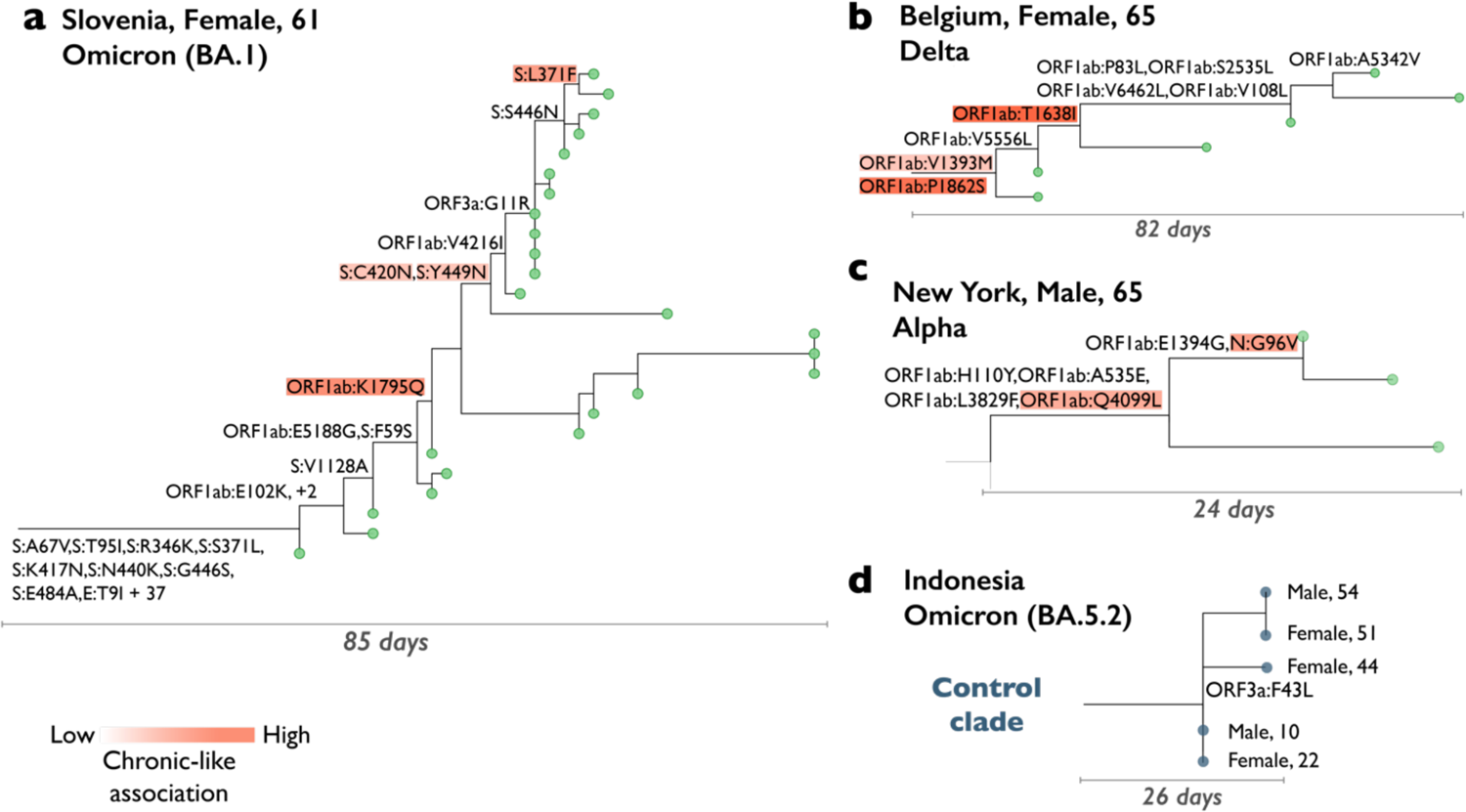
Examples of chronic-like clades detected by our approach. Panels a-c depict a chronic-like clades, where all sequences (tips) were presumably sampled from the same individual. Panel d illustrates a control clade. Only non-synonymous mutations are shown along the branches. In panels a-c, mutations are color-coded based on their association with the group of chronic-like clades, with stronger shades of orange indicating a stronger association (see main text below on language models).

We next went on to examine the evolutionary divergence in the chronic-like clades as compared to the controls (Methods). Notably, this is a challenging endeavor as the clades we analyzed span a very short time (ranging from 21 to 241 days), and sequences are not independent due to shared ancestry. Moreover, sequences towards the tips of the tree tend to be enriched with slightly deleterious mutations (so called incomplete purifying selection) ^24^. Nevertheless, given the similar distributions of sizes and times across clades and controls (Fig. S1), we considered that averaging the regression slopes across a set of clades may allow us to compare trends between different sets of clades. We thus employed a simple linear regression and regressed the number of mutations from the ancestral node, against the calendar day, for all sequences in each clade. The average slope of the regression lines was significantly higher in chronic-like clades as compared to controls (*p* < 10^-4^, t-test), with average slopes corresponding to 16.63 and 12.56 mutations per year, respectively (Table S3). Reassuringly, the average slope in controls was in line with estimates of divergence obtained previously across large sets of sequences from global transmission ^22^. When focusing on synonymous mutations, we did not find any significant differences between the slopes of chronic-like versus control clades (*p* = 0.37, t-test). However, the average slope of non-synonymous mutations was significantly elevated, with a value of 13.8 in the chronic-like clades compared to 7.7 for control clades (*p* < 10^-10^, t-test). Overall, these results are highly suggestive of adaptive evolution occurring in the chronic-like clades.

We examined whether a specific VOC background was associated with different patterns of evolution in the chronic-like clades. An examination of the Sackin index did not reveal any differences among the chronic-like clades from different variants (Fig. S2). However, we noted that the regression slopes for non-synonymous mutations were lower in Delta as compared to BA.1 chronic-like clades (*p* < 0.05; one way ANOVA with Tukey’s multiple comparisons test). This could not be explained by differences in clade size or span of sampling dates (Fig. S1). We could not conclusively determine if a particular genomic region is driving the differences between Delta and BA.1 (Table S3) (see discussion).

Next, we tested whether mutations associated with chronic-like clades can “predict” mutations in future variants, i.e., variants that emerged after the dates when the chronic-like clade was sampled (see Fig. 3a). We focused on a set of eleven positions in the spike receptor binding domain (RBD) where rampant convergent evolution was detected since the emergence of BA.2 ^25^, most likely signifying they are adaptive mutations (Fig. 3b, Fig. S3). Strikingly, ten of these eleven positions were mutated in chronic-like clades that preceded the date when these mutations began increasing in frequency (with the exception of S:L452Q, Fig. S3). In BA.1 chronic-like clades >70% of the mutations were detected whereas in BA.2 (where our sampling was more limited) almost 30% of the mutations were detected (Fig. 3c).

**Figure 3.**
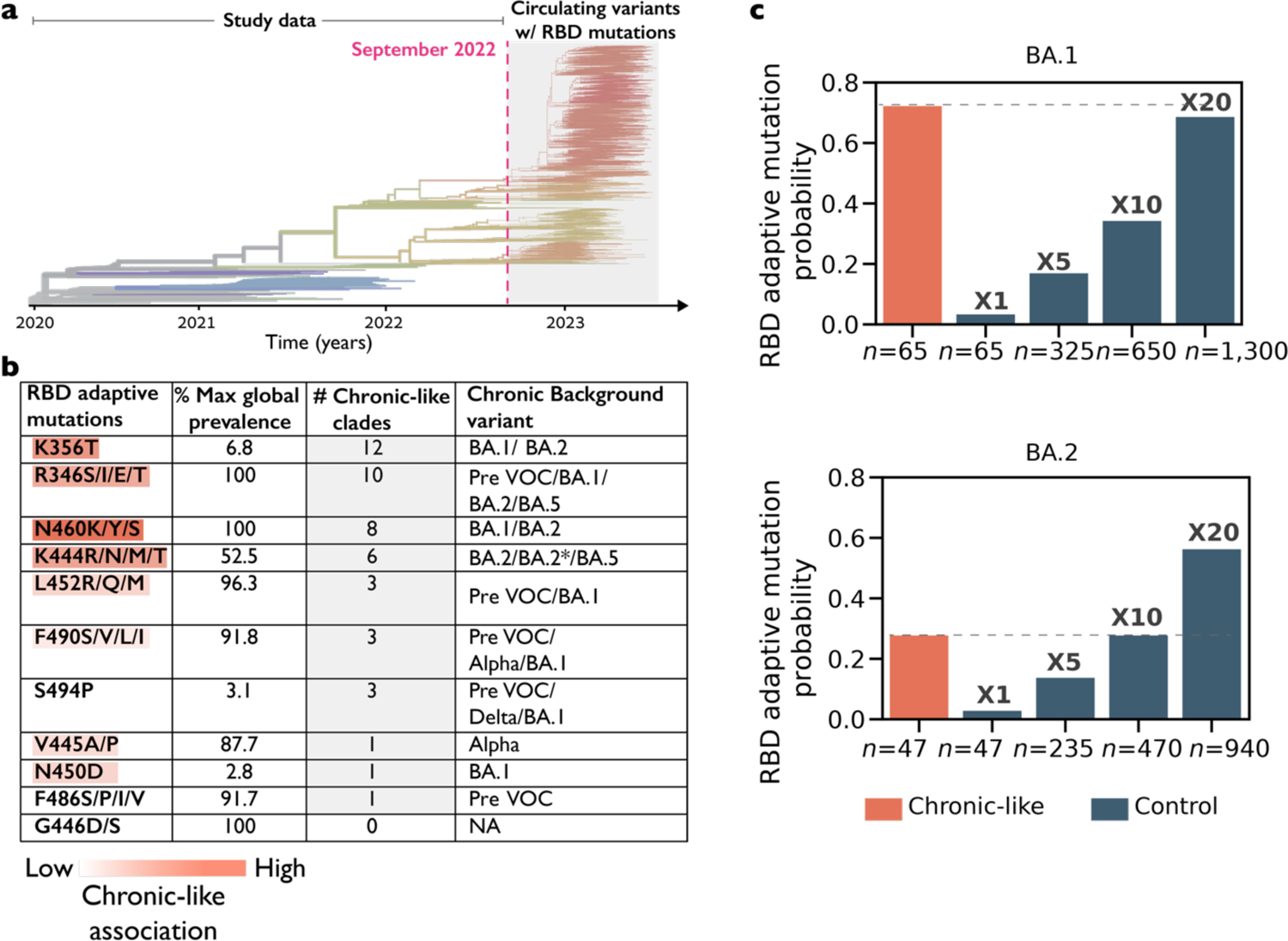
Adaptive mutations forecasted in chronic-like clades. (a) Timeline of the study data indicating the last date of sampling (dashed) line, projected onto a phylogeny of SARS-CoV-2 derived from nextstrain.org ^3^ (b) A list of RBD positions where highly convergent mutations occurred in sub-lineages of Omicron circulating since the summer or fall of 2022. Ten of eleven of these positions were mutated in chronic-like clades. Mutations are color-coded based on their maximal association with the group of chronic-like clades, with stronger shades of orange indicating a stronger association (see main text below on language models). Max global prevalence refers to the maximal proportion of sequences across all global sequences, where this mutation was observed, at any time point in the pandemic (Fig. S3) (b) The probability of detecting RBD adaptive mutations in chronic clades compared to their probability of detection in increasing sample sizes of control clades.

Overall, our results suggest that chronic infections speed up the probability and hence the rate of adaptive evolution. Across balanced sets of equally sized clades, there were more non-synonymous mutations in chronic-like clades, particularly in the S1 domain of spike and in the RBD (Fig. 1d). Accordingly, it is possible that increasing the viral population size in the community, which is akin to more transmission chains, would lead to an increased probability of observed beneficial mutations. To test this, we increased the sample size of controls and tested the fraction of control clades where we observe the list of convergent spike mutations in Fig 3b. We found that a 10-20-fold increase in the size of the controls led to equal probabilities of observing these adaptive RBD mutations between controls and chronic-like clades (Fig. 3c).

We wished to predict the sets of mutations that are most associated with chronic-like clades and use the associations to predict new clades when reliable metadata was absent. However, performing these tasks presented several challenges: The phylogenetic-based analysis was both computationally intense and suffered from inaccuracies in the phylogenetic tree. Further, only about 25% of sequences bore reliable metadata; most sequences were labeled as “unknown” in at least one of the categories sex or age. These challenges led us to adopt a deep learning approach that relied on language models suitable to deal with large, noisy, and unlabeled data and also allowing us to minimize our dependency on the phylogeny.

In our language model, “words” are mutations compared to the reference genome sequence, and thus a given SARS-CoV-2 genome is a sequence of words representing all its mutations as compared to a reference genome (Methods). We began by pre-training a BERT masked language model ^26^ on the set of 11.7 million sequences after excluding cases, controls, and other low-information sequences (Methods). We filtered out very low frequency mutations (Methods), so that overall, our model consisted of a vocabulary of 38,000 unique mutations (formally called tokens). We learned the conditional probabilities associated with observing a specific mutation within various mutational contexts. This allowed us to account for differences stemming from the background variants. Consequently, we reasoned that the probability of a given mutation emerging could vary depending on the VOC background and timing (pre- or post-vaccination/convalescence). This is in part due to epistatic interactions among mutations, as observed repeatedly for SARS-CoV-2, particularly in Omicron ^27–29^.

Our language model provided a numerical representation for each mutation. Projecting these representations onto a two-dimensional space allowed us to verify that the model effectively separated sequences into their respective Nextstrain clades (Fig. S4, Methods), reassuring that the model captures similarities among mutational profiles. We next fine-tuned our pre-trained model to perform a classification task of distinguishing cases from controls. For the classification task, we employed the BERT model with two additional neural network layers dedicated to classification. To test our classification process, we adopted a sequential cross-validation approach based on the time of emergence and the background variant: the model was trained iteratively, where data from preceding variants was used as training data for each new variant tested. We found that the classifiers were successful in distinguishing between cases and controls with an average Area Under the Precision-Recall curve (AUPR) of 0.88, weighted by clade size and the number of mutations in the clade (Methods, Fig. S5).

To gain further insights into the classifier’s predictions for the cases, we employed LIME (Local Interpretable Model-agnostic Explanations) ^33^, a technique used to identify the “words” that have the greatest impact on model predictions and the reliability of each such inference (Methods). We focused on LIME scores higher than 0.05, which correspond to the 75% quantile. Several interesting observations emerged from this analysis. As expected, for most variants, non-synonymous spike mutations were most predictive of belonging to a chronic-like clade (Fig. 4a). Six of the eleven convergent RBD positions noted above (Fig. 3) had high LIME scores (and an additional two had LIME scores slightly below the 0.05 cutoff). Out of a total of thirteen positions with high LIME scores in BA.1/BA.2 chronic-like clades (Supplementary Dataset 6), six indeed were mutated in currently circulating strains (Fig. 3).

**Figure 4.**
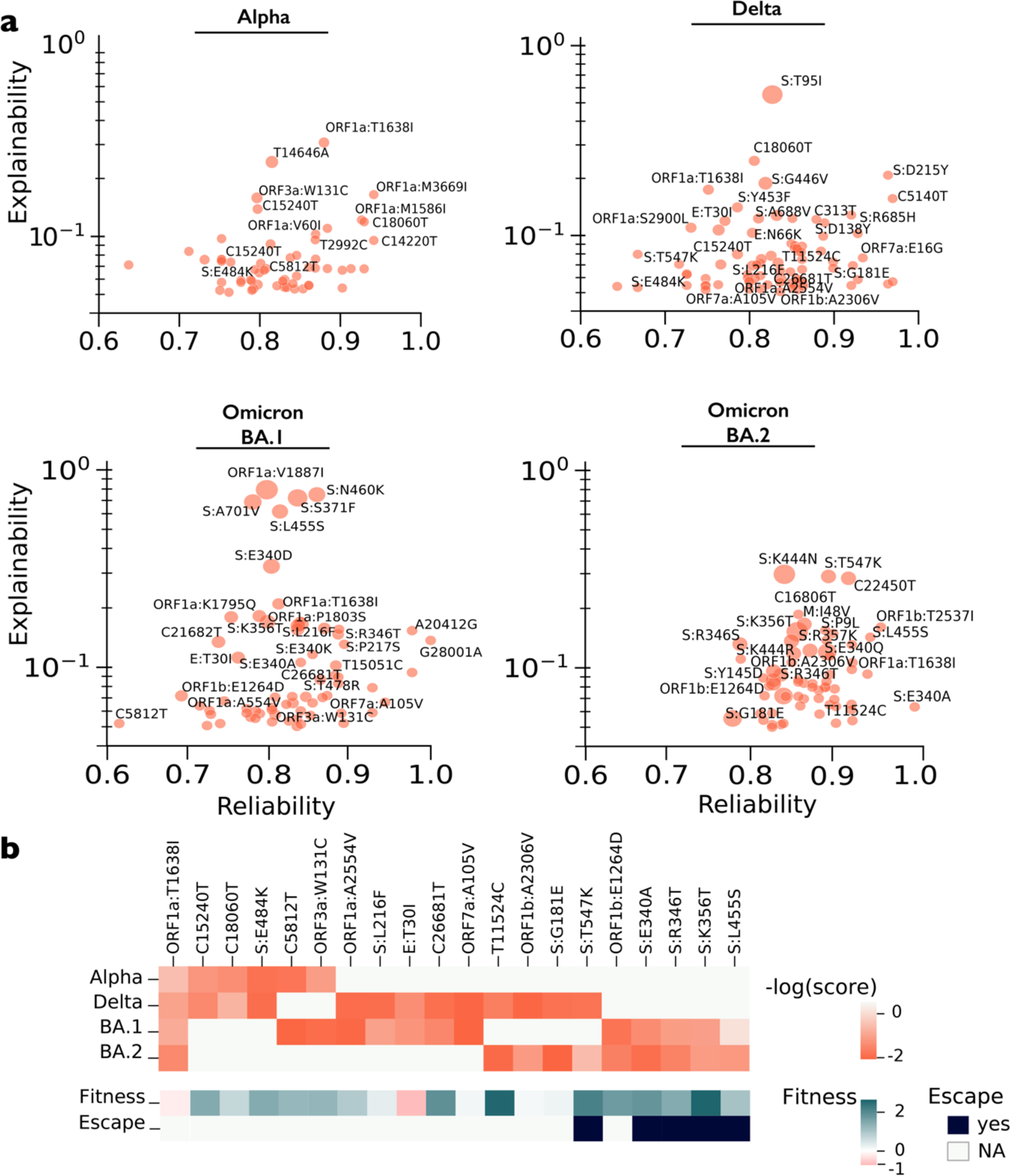
Mutations associated with chronic-like clades. (a) Mutations that best explain model predictions of chronic-like clade. Each panel depicts the mutations per a given background variant, with only those with an explainability score higher than 0.05 (75^th^ quantile) shown (y-axis). The x-axis represents the average LIME R^2^ score, indicating the prediction reliability. Dot size correlates with the number of samples in which the word was observed. Only mutations with an explainability score higher than 0.1 are labelled, as well as those with a score higher than 0.05 that recur across different background variants (panel b). The full list of mutations is available in Supplementary Dataset 6. (b) Recurrent mutations associated with chronic-like clades, across different background variants. All mutations with an explainability score higher than 0.05 that appeared in at least two variants were used for the intersection. Fitness is based on inferences derived from mutations abundance across all globally circulating sequences ^30^ and antibody escape is based on inferences from deep-mutational scans using a variety of different types of antibodies ^31, 32^. (a,b) Synonymous nucleotide substitutions are denoted by their genomic position, whereas amino-acid replacements are denoted by the protein name and amino-acid replacement.

We noted that different variants were associated with different mutations. For example, S:E484K that we originally observed in many WT chronic infections, was associated with Alpha and Delta, but not with Omicron backgrounds, where an S:E484A had been fixed as a lineage-defining mutation. While only a few synonymous mutations were detected by LIME, interestingly, these were associated with many different background variants (Fig. 4b). Moreover, all shared spike RBD mutations were associated with antibody evasion, which is expected. Finally, we noted that a recent analysis has shown that a large proportion of mutations in SARS-CoV-2 are under purifying selection and are predicted to have low fitness ^30^. On the other hand, all but two of the shared mutations in our chronic-like clades were predicted to have high fitness. Two mutations (ORF1a:T1638I, E:T30I) were predicted to have negative fitness (Fig. 4b) (see discussion).

We went on use the model to predict chronic-like infections from sequences with missing metadata, in the absence of a phylogeny. To this end, we used the “words” (i.e., mutations) that we previously found as most strongly associated with controls or chronic-like clades and searched for clades with the highest probability of being derived from chronic infections. The number of predictions naturally depends on the prediction score cutoff we define (Fig. 5a).

**Figure 5.**
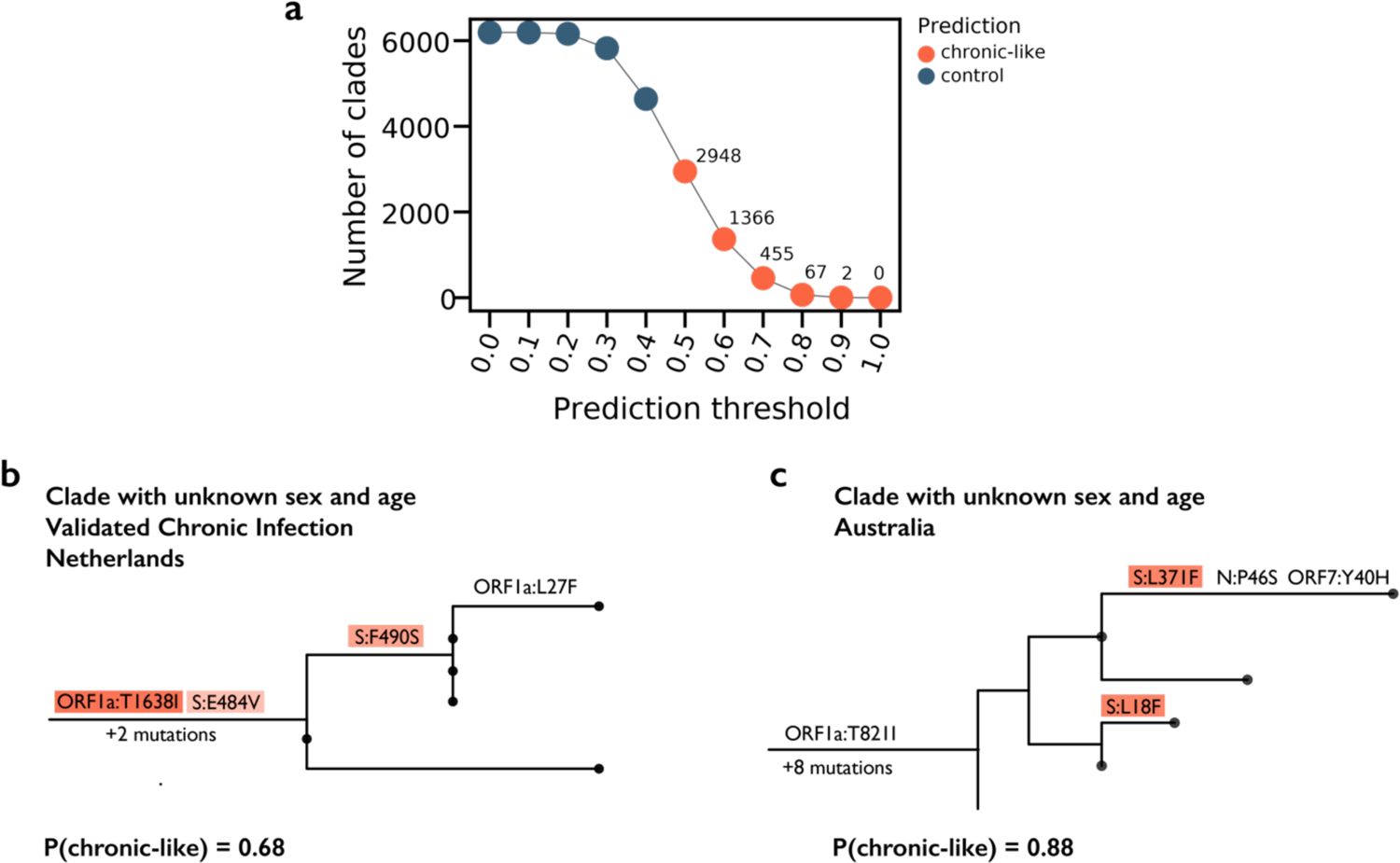
Prediction of chronic-like infections in the absence of metadata and phylogeny. (a) The number of chronic-like clades inferred given different prediction thresholds. (b-c) Two examples of clades inferred as chronic-like infections, with mutations driving the prediction color-coded as in Fig. 2.

Given that we detected 271 chronic-like clades from 25% of the data that had reliable metadata, naively we expected three times more such clades (∼800) in the remaining 75%. A prediction score cutoff between 0.6 and 0.7 resulted in such an estimate. We present two examples of clades inferred with a prediction score higher than 0.65 in Figs. 5b,c. One of these cases is actually a *bona fide* chronic infection where sequences were submitted with missing metadata. This case was identified by directly contacting the authors ^34^ (Fig. 5c; Methods), and the second is a completely novel prediction.

## Discussion

In this work, we harnessed a very large volume of SARS-CoV-2 sequencing data and metadata to search for sequences that are likely derived from chronic infections. Our approach relies on the idea that there is deposition of viral sequences and metadata from de-identified individuals. In some cases, repeated sequencing is performed from the same individual who may be chronically infected. We search for signatures of such chronic infections by first directly mining a huge phylogenetic tree and next by using deep learning approaches that rely on our first findings. We go on to show that the chronic-like clades we find, particularly the most recent ones, are predictive of mutations that are later part of circulating strains.

There are some important limitations to the approach we put forth. First, there is no realistic way that we can verify each of our inferences of chronic-like clades. However, the analyses we show herein suggest that this set of clades is strongly enriched for chronic infections. Second, we most likely dramatically underestimate the number of chronic-like clades: we do not account for chronic infections sampled less than three times, we do not account for chronic infections where sequence data were not of high quality, and we do not account for chronic infections that lead to onwards transmission. A recent paper has estimated that less than 3% of chronic infections lead to onwards transmission ^15^, suggesting the latter factor is minor; however, this does mean we might be overlooking chronic infections with features that allow such onwards transmission to occur.

We interpret the results we found with regards to chronic-like clades. Interestingly, we found a lower rate of non-synonymous divergence in Delta as compared to BA.1. We could not conclusively pinpoint which gene or genomic region was responsible for this difference; we noted a borderline insignificant p-value for the spike gene (*p*=0.1, ANOVA, *p*=0.06, Tukey test; Table S3). Moreover, it was not clear whether the difference was driven by a higher rate of adaptive evolution in BA.1 chronic infections, possibly due to increased use of monoclonal antibodies, or due to an inherently lower rate of adaptive evolution in Delta chronic infections. The latter would be interesting, as Delta is the only VOC that lacks the N501Y mutation, which has been shown to form epistatic interactions that enable multiple additional antibody escape mutations ^27–29^. Most of our Delta chronic-like clades were between May 2021 and March 2022, when vaccination and convalescence rates varied widely across the globe, and this may have impacted the probability of antibody escape mutations. However, we do not have enough data to shed more light on this finding.

We go on to discuss the features of the mutations most associated with chronic infections (Fig. 4a). The strongest signal was most often derived from non-synonymous spike mutations that have been shown to promote ACE2 binding and/or antibody evasion, but was also driven by mutations in other genes, predicted to be high fitness mutations (Fig. 4b). Of particular note are two mutations (ORF1a:T1638I, E:T30I) that were highly associated with chronic-like clades yet were inferred to have negative fitness, suggesting they are generally absent in globally circulating lineages ^30^. It is likely that these mutations endow a fitness advantage during intra-host evolution yet a disadvantage during inter-host transmission. E:T30I has been previously noted ^5,14^ but its role in inter-host evolution is not yet understood. ORF1a:T1638I (nsp3:T820I) is the mutation most commonly associated with all backgrounds, and was found to be associated with HLA-A∗01:01-allele restricted escape from CD8+ T-cells ^35^. If so, it is possible that it may be advantageous on the background of certain HLA backgrounds but may be disadvantageous on other backgrounds, which could explain the absence of this mutation in the global phylogeny. Alternatively, it may have a different negative impact on transmission.

Our results highlight that adaptive evolution is enhanced during chronic infections as compared to evolution along transmission chains. This may be due to several non-mutually exclusive reasons: higher selective pressures, large viral population size, and the ability to cross fitness valleys through epistatic interactions among mutations ^36^. In the initial stages of the pandemic, it is likely that selective pressures differed more dramatically between acute and chronic infections. In chronic infections, the weakly functioning adaptive immune pressure, coupled with antibody-based treatments, would have created strong pressure for the virus to evade immunity. However, before vaccinations and widespread infections, most acute infections were of immunologically naïve individuals and would not have elicited strong immune selection pressure. This however changed dramatically over time, as rates of vaccination and convalescence (leading to so-called hybrid immunity) increased dramatically. Thus, today it is possible that the immune pressures the virus experiences in acute versus chronic infections are more similar. On the other hand, there is still a dramatic difference in the viral population size between acute and chronic infections. During acute infection and subsequent transmission, viruses undergo a dramatic population bottleneck, estimated at *n*_b_=1 ^11, 12, 37^ This would dramatically dampen the rate of adaptive evolution as compared to chronic infections where the virus population size is much larger.

During late 2022 and 2023, many new sub-variants that emerged (e.g., lineages deriving from XBB or BA.2.75) were not highly divergent but were rather characterized by more gradual accumulation of mutations. It is possible that this was enabled by the explosive number of transmissions during the Omicron waves, that increased the fixation probability of adaptive mutations. Thus, when global transmission was relatively limited and when population immunity was low, chronic infections served as an important evolutionary catalyzer that allowed rare events to occur. Later on, as transmission increased, coupled with increasing hybrid immunity in the population, we surmise that evolution along transmission chains allowed enough opportunities for adaptive point mutations to fix. Of note, it is still possible that a future chronic infection will yield a combination of mutations that is less likely along transmission chains (epistatic interactions). Alternatively, we may be at a situation that is very similar to Influenza evolution, where point mutations at viral epitopes endow a large fitness advantage ^38^.

The analysis of sequencing data often heavily relies on phylogenetic methods and models that are quite computationally expensive. As the volume of biological sequence data continues to increase, language models are emerging as valuable tools ^18, 39, 40^. We have put forward the idea of using language models and deep learning to infer chronic infections and show as a proof-of-concept that this approach can most likely augment our inferences. In an era of ever-growing pathogen sequencing, we envisage that monitoring chronic-like clades using the approaches described herein may be valuable to predict the next mutations and the next variant that may take-over global transmission.

## Methods

### Sequence dataset curation and pre-processing

We accessed the GISAID database ^41–43^ on September 17^th^, 2022 and downloaded a total of 13,165,623 SARS-COV-2 sequences with their associated metadata. We began by employing stringent quality control and used an initial filtering step based on Nextclade version 2.5 ^44^, which scores sequences based on a series of comprehensive criteria. We used all Nextclade defaults but changed the mixed sites threshold to 30. We also excluded from these criteria the “private mutations” criterion, since we considered private mutations may very well characterize chronic infections. Sequences with a final quality score of “bad” or “mediocre” were removed from our analysis, while only those labeled as “good” were retained. Additionally, we removed sequences with ambiguous or conflicting dates (missing/partial dates, or a submission date earlier than collection date). We manually curated the age and sex metadata fields, resolving conflicts resulting from reporting in languages other than English. We considered ages below one year as one year old and maintained age ranges as provided (e.g., two samples labeled as age 10-19 were considered as samples of the same age). Samples where metadata was missing or corrupted were labeled with “unknown” in the relevant field. Overall, these steps resulted in a reduction of approximately 11% of the original number of sequences and yielded a final dataset of 11,717,404 sequences.

### Phylogeny-based inference of chronic-like clades

We used a global SARS-CoV-2 phylogenetic tree that is constantly updated with sequences added to the GISAID database. The tree was reconstructed using the UShER algorithm ^45^ and was kindly provided by Angie Hinrichs on August 25^th^, 2022. The mutational path of each sequences is also included with the UShER tree, and includes for each sequences the step-by-step mutations that occurred from the root of the tree till each leaf, based on ancestral sequence reconstruction ^45^. We used this tree to identify “chronic-like” clades, defined as clades potentially derived from an individual with a chronic SARS-COV-2 infection:

Formally, we define a “chronic-like” group *M* in the tree *T* as a group of sequences {*S*_1_, …, *S*_n_} ∈ *T* that meets the following criteria: *M* defines a monophyletic clade with at least *n* = 3 leaves but no more than 40 leaves; all *S*_i_ ∈ *M* share the precise same location, the sequences {*S*_1_, …, *S*_*n*_} span at least 21 days, and 75% of the sequences in *M* share the same age and sex (excluding “unknown” samples). Notably, we relaxed the last assumption from 100% to 75%, after extensive manual testing that revealed that in many cases sporadic sequences were included in the clade either erroneously or correctly but with missing metadata. We recovered sequences with ambiguous dates that were excluded in the initial quality control if they were part of a candidate clade (e.g., a sequence excluded since it was reported as February 2021 and was re-included if the chronic-like clade spanned this month).

Given the high proportion of data with an “unknown” label, we extracted additional clades to be further prioritized as “chronic-like” groups. We define an unknown group *U* in the tree *T* in the same manner as defined above, except that both age and sex are unknown. This yielded 18,760 clades.

### Control clades

Finally, we define a control group *C* in the tree *T* as a group *C* = {*S*_1_, …, *S*_*n*_} ∈ *T* that meets the following criteria: 3 ≤ *k* ≤ 40, all *S*_i_ ∈ *C* share the same location, the sequences {*S*_1_, …, *S*_*n*_} span at least 21 days, yet we ensured that the ages and sexes differ, i.e., at least four different combinations of age and sex exist in the group. This yielded 15,163 clades.

Due to the large difference in sample size between the cases and controls, we used bootstrapping to control for sample size. Specifically, we performed stratified sampling with replacement of *n* = 10^%^ subgroups from the control group, each sized 271 (identical to the number of chronic-like clades), stratified by the Nextclade clade to allow for similar background variant distribution. This bootstrapping approach allowed us to calculate average values for the control clades across different measures described below.

### Bona fide chronic infections

We relied on our previous publication that includes *n* = 27 chronic infections ^5,34^ and added on sequencing data from^34^ of *n* = 5 Omicron chronic infections (data was obtained by directly contacting the authors).

### Assignment of mutations to clades

We set out to find the within-clade evolution of each of the chronic-like clades that we inferred. Notably, the UShER mutational path reports nucleotide mutations and lacks indels whereas the Nextclade annotation includes amino-acid replacements and indels. Therefore, for each clade (chronic-like or control), we first extracted the set of all mutations from the sequences using Nextclade mapping ^44^. Then, we intersected these data with the UShER mutational path and removed from this set all mutations that occurred up to the ancestral node of each chronic-like clade. We included the mutations on the branch leading to this. At this stage we also excluded indels from this analysis since they were not included in the UShER mutational paths. Of note, all mutations in this manuscript are reported with respect to the ancestral Wuhan-Hu-1 reference genome sequence (GenBank ID NC_045512).

### Position masking

We masked all lineage defining mutations from the analysis since we noted that some sequences were erroneously assigned with the reference sequence nucleotide, presumably when sequencing coverage was low or when sequencing quality was poor in a given region. To this end each sequence was assigned a clade based on Nextclade, and lineage-defining mutations were based on https://github.com/neherlab/SARS-COV-2_variant_rates/blob/f807ae8015f6004dec38e6c757d5cedb9625cd6e/data/clade_gts.json#L4^22^. Additionally, we masked mutations flagged as problematic positions in this table (https://github.com/W-L/ProblematicSites_SARS-CoV2/blob/master/problematic_sites_sarsCov2.vcf) ^46^. Finally, we masked the two positions upstream and downstream of each masked position described above.

### Binned distributions

To compare the distribution of mutational counts between the chronic-like clades and controls, we divided the entire genome into 500-position long bins, denoted as *bin*_i_, where *bin*_i_includes all mutations observed in positions [*i* ⋅ 500, *i* ⋅ 500 + 500). We used a binomial test to identify bins significantly enriched for mutations.

### Sackin index

We extracted the sub-tree for each clade using the UShER platform ^45^, and calculated the Sackin index ^47^ as a measure of tree imbalance using Python’s dendroPy framework^48^.

### Linear regression for within clade mutation accumulation rate

We used an ordinary least-square (OLS) linear regression model to assess the slop of each chronic-like and control clades. For a given clade, we obtained for each sequence the number of within-clade mutations since the regression number of mutations per sequence against date. OLS was performed using the python statsmodels version 0.13.5 ^49^.

### Language model for mutation representation

To create a corpus of all mutations in the data, we used the set of sequences described above and constructed “sentences” comprised of “words” (tokens), each representing a mutation relative to the reference sequence. We included only mutations in coding regions and used the mutation annotation by Nextclade. Non-synonymous mutations were represented by the gene where they occurred and the associated amino acid replacement (e.g., S:D614G), while synonymous mutations were represented by the genome location and associated nucleotide change (e.g., C3067T). Deletions and insertions were also included and were represented as obtained by NextClade (e.g., 27871 for a deletion, 20:GGA for an insertion) ^44^. For each sample in the GISAID dataset, a sentence was constructed to encompass all the mutations present in that sample. These mutations were sorted based on their genomic location to ensure a coherent representation within the sentence.

We limited the model vocabulary to mutations that appeared at least 45 times in the dataset, to allow for a reasonable vocabulary size of 38,000 unique tokens. Next, we trained a BERT model ^26^ from scratch on the task of masked language modeling to generate a numerical representation for each sample in the dataset. We excluded sequences that were selected as chronic-like clades, control clades, and clades with unknown metadata that were used later for classification and prediction. This resulted in a dataset of 10,646,407 sequences. We used 90% of the data for training and the remaining 10% for validation.

For tokenization, we applied a custom BERT tokenizer from the Hugging Face library ^50^ that splits sentences into tokens based on whitespace. We set the sentence length limits to be between 5 and 160 words. During training, the masking probability was set to 0.2, with a per-device training batch size of 10 and a validation batch size of 64. We used gradient accumulation steps of 8, which resulted in the final training batch size of 640 and a validation batch size of 512. The model was trained with the default optimizer AdamW ^51^ for two epochs with an initial learning rate of 1e-5. The best model was chosen based on the minimal evaluation set loss. The model was trained on a single NVIDIA RTX A6000 GPU with 48G RAM and 8 CPUs, taking a total of 48 hours.

### Mode fine-tuning for chronic-clades classification

We used our pre-trained BERT model to classify chronic-like clades versus controls. Each clade was represented by a sentence that included all of the clade’s corresponding mutations.

Specifically, clade *c*_i_ was represented by the sentence *S*_i_ = {“*m*_1_ *m*_2_ … *m*_n_” |*m*_j_ ∈ *S*(*C*_i_)}, where *m*_j_ is a mutation (token) and *S*(*C*_i_) denotes all sequences within clade *C*_i_. To ensure consistency, mutations were sorted based on their genomic location.

To prevent potential confusion between lineage-defining mutations that occurred in the past and their subsequent appearance in chronic clades, all lineage-defining mutations according to the Nextclade variant were removed. Masking of problematic positions/mutations was performed as described above.

The classification process was performed based on VOC chronology, utilizing three distinct folds. The dataset was divided into five main groups: pre-VOC, Alpha, Delta, BA.1 and BA.2, along with the remaining variants. For the non-pre-VOC variants, we considered the preceding variants for training and tested on the relevant variant (Table S1).

To ensure a balanced representation of the control data, a down-sampling technique was applied. Initially, the control clades were filtered based on the Sackin index, including only those with a value lower than 1.44. This threshold was determined by averaging the values obtained through bootstrapping analysis. The objective was to intentionally select control clades who are most likely derived from transmission chains and unlikely to have derived from a chronic infection. Following the initial sampling, we performed an additional round of down-sampling down to n=∼270 control clades, to ensure that the Nextstrain background variants and clade size are balanced across the set of controls and chronic-like clades.

To optimize the classification process given the relatively small sample size, we modified the BERT for Classification architecture obtained from Hugging Face. Specifically, we froze all embedding and encoder layers, allowing changes only to the last layers responsible for pooling and classification. The training procedure encompassed 30 epochs, and for each fold, the model with the lower evaluation loss was selected (Fig. S4). Throughout the training phase, a batch size of 64 was employed, while a batch size of 32 was used for the evaluation set. To assess the performance of each model, we utilized two metrics: ROC AUC and AUPR (weighted by clade size and the number of mutations).

### Model Explainability

Local Interpretable Model-Agnostic Explanations (LIME) ^33^ was applied to understand the underlying reasoning behind how the classifier inferred chronic-like clade. In each test fold (Alpha, Delta, BA.1,BA.2), we identified the top 10 mutations with the highest LIME scores for each clade. These scores were aggregated by introducing a mutation score per background variant, denoted as *S*_*v*_(*m*), which sums the LIME scores across all samples within variant *v*’s test set. This aggregation approach enhances the significance of mutations that consistently appear across multiple samples, reinforcing their predictive potential. Additionally, as LIME fits a regression line per clade in order to generate inferences, the *R*^2^ serves as a measure of result reliability, and here it was was averaged across clades.

### Data availability

All data supporting this study’s findings will be deposited to the Zenodo database and assigned a permanent DOI; meanwhile, they are available here: https://tinyurl.com/4jxuvbak. These data include the chronic-like and control clades identifiers, mutations, and sub-trees. They also include language model raw corpus files, trained models, and all predictive mutations for the chronic-like clades groups. The full description is available in the Supplementary information and the attached Readme file in the link provided.

### Code availability

All models used for training and scripts used for analysis are available in the repository https://github.com/Stern-Lab/chronic-covid-mlm.

## Acknowledgments

We would like to thank Pleuni Pennings, Jesse D. Bloom, Avigdor Eldar, Daniel Weissman and Angie S. Hinrichs for helpful discussions and comments on the manuscript. We thank all the sequence contributors to GISAID who made this work possible. This study was supported by grants to AS: an ERC starting grant 852223 (RNAVirFitness), an Israeli Science Foundation grant 1930/22, and a grant from the Tel Aviv University Center for Combatting Pandemics. This study was also supported by fellowships to S.H. and D.M. from the Edmond J. Safra Center for Bioinformatics at Tel Aviv University and a Fellowship to D.M. from the Azrieli Foundation.

## Contributions

Conceptualization: S.F., A.S., D.M. and S.H. Phylogenetics and bioinformatics: S.H. and D.M. Initial proof of concept analyses: S.F. Language model design and implementation: D.M. and D.B. Supervision: D.B and A.S. Writing: S.H., D.M, A.S. All authors read and revised the paper.

## Notes

https://tinyurl.com/4jxuvbak

### Competing Interest Statement

The authors have declared no competing interest.

