## Supplementary information for "Using big sequencing data to identify chronic SARS-Coronavirus-2 infections"

### Using big sequencing data to identify chronic SARS-Coronavirus-2 infections: a window into future evolution

This supplementary information contains figures, tables and supplementary datasets related to the publication. All Supplementary Datasets described will be deposited to the Zenodo database and assigned a permanent DOI; meanwhile, they are available here: https://tinyurl.com/4jxuvbak

##### Supplementary Tables

Supplementary Table 1 - Variant definition using NextStrain clades
Supplementary Table 2 - Country distribution across the 271 chronic-like clades divided by continent
Supplementary Table 3 – Slopes for regression lines of mutations against sampling dates

##### Supplementary Figures

Supplementary Figure 1 - Per-variant clade size and time interval distributions
Supplementary Figure 2 - Distribution of Sackin index across 271 chronic-like clades
Supplementary Figure 3 – Global prevalence of RBD convergent mutations across time
Supplementary Figure 4 - Language model Embedding representation and training
Supplementary Figure 5 – Classification performance assessment

##### Supplementary Datasets

Supplementary Dataset 1 – Chronic-like samples data and identifiers
Supplementary Dataset 2 – Control samples data and identifiers
Supplementary Dataset 3 – USHER down sampled phylogenetic trees and Sackin index measurements
Supplementary Dataset 4 – Examples of chronic like clades presented in Figure 2 and 4
Supplementary Dataset 5 – Regression evolutionary clade rates
Supplementary Dataset 6:

1. Language model corpus
2. Chronic-like clades and controls sentences for classification
3. Model performance tables
4. LIME explainability results tables
5. Clades with unknown data predictions

**Supplementary Table 1.** Variant definition using NextStrain clades[1,2] .

| Variant | NextStrain clade |
|---|---|
| Pre VOC | 19A,19B,20A,20B,20C,20E,20G,20D |
| Alpha | 20I |
| Delta | 21J,21I |
| Omicron | 21K,21L,22A,22B,22C |
| Other | 20H,20J,21F,21B |

**Supplementary Table 2.** Country distribution across the 271 chronic-like clades divided by continent.

| Continent | Countery | Chronic-like clades |
|---|---|---|
| Africa | South Africa | 2 |
| Asia | Indonesia | 1 |
| | Japan | 2 |
| | Israel | 5 |
| | India | 13 |
| Europe | Croatia | 1 |
| | Slovakia | 1 |
| | North Macedonia | 1 |
| | Austria | 3 |
| | Romania | 3 |
| | Netherlands | 3 |
| | Ireland | 3 |
| | Luxembourg | 7 |
| | Belgium | 8 |
| | Germany | 11 |
| | Sweden | 14 |
| | Italy | 24 |
| | Slovenia | 30 |
| | Spain | 33 |
| | France | 48 |
| North America | Mexico | 2 |
| | Canada | 35 |
| | United States of America | 55 |
| Oceania | Australia | 7 |
| South America | Argentina | 1 |
| | Peru | 1 |
| | Brazil | 5 |

**Supplementary Table 3.**

| Variant | Label | Overall[1] | Synonymous[2] | Non-synonymous[3] | Spike non-synonymous[4] |
|---|---|---|---|---|---|
| Alpha | control | 12.50 | 6.11 | 7.62 | 3.01 |
| | chronic-like | 16.32 | 6.72 | 13.02 | 8.06 |
| Delta | control | 14.67 | 6.88 | 8.96 | 2.17 |
| | chronic-like | 10.38 | 4.48 | 7.98 | 4.88 |
| BA.1 | control | 10.65 | 5.20 | 6.95 | 2.63 |
| | chronic-like | 22.51 | 5.97 | 18.97 | 10.25 |
| BA.2 | control | 10.90 | 5.66 | 6.58 | 2.90 |
| | chronic-like | 15.55 | 5.087 | 13.02 | 8.69 |

1 ANOVA: p =0.05, Tukey: Delta-BA.1 p=0.03, all other pairs are non significant
2 ANOVA: p =0.87, Tukey: all pairs are non significant
3 ANOVA: p =0.019, Tukey: Delta-BA.1 p=0.013, all other pairs are non significant
4 ANOVA: p =0.10, Tukey: all pairs are non significant
* ORF1a and ORF1b ANOVAs are non significant for any group presented.

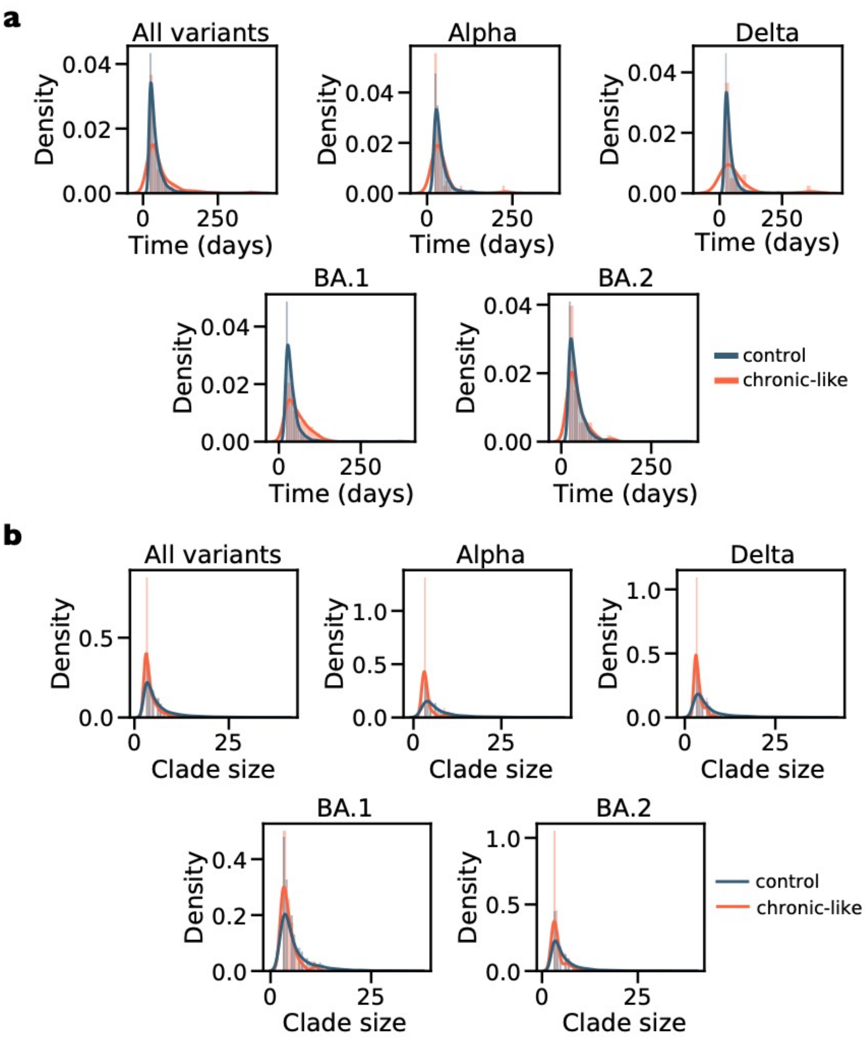


**Supplementary Figure 1. Per-variant clade size and time interval distributions separated by class.** (a) Time interval distribution in days. (b) Clade size distribution.

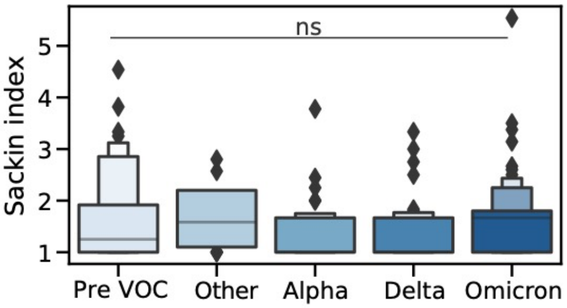


**Supplementary Figure 2. Distribution of Sackin index across 271 chronic-like clades, categorized by Nextstrain variant**. Variants were classified based on Table S2. An ANOVA test was conducted to assess differences among the variant groups, revealing no statistically significant differences ($p = 0.16$).

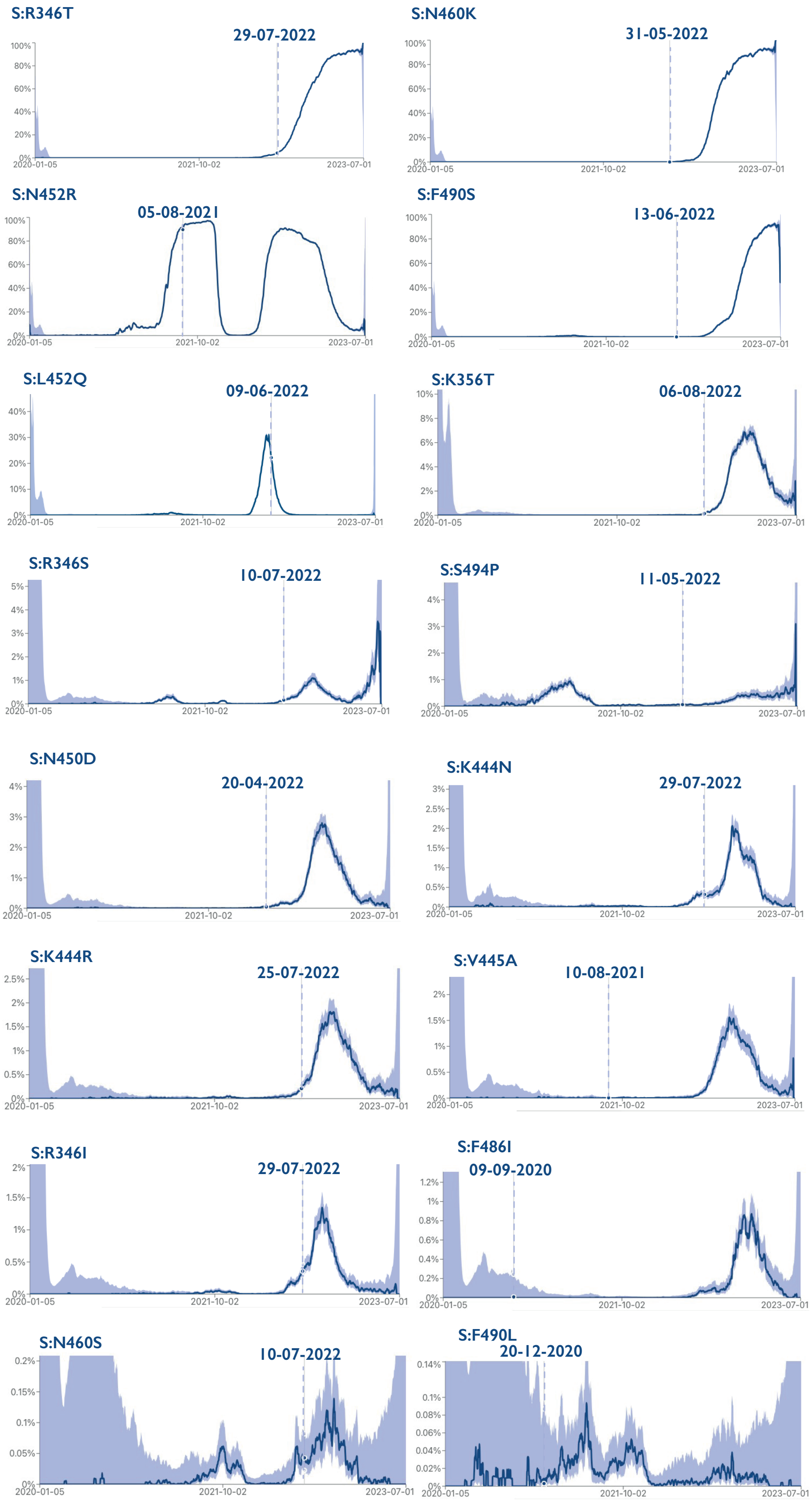
S:R346T
29-07-2022
S:N460K
31-05-2022
S:N452R
05-08-2021
S:F490S
13-06-2022
S:L452Q
09-06-2022
S:K356T
06-08-2022
S:R346S
10-07-2022
S:S494P
11-05-2022
S:N450D
20-04-2022
S:K444N
29-07-2022
S:K444R
25-07-2022
S:V445A
10-08-2021
S:R346I
29-07-2022
S:F486I
09-09-2020
S:N460S
10-07-2022
S:F490L
20-12-2020
2020-01-05
2021-10-02
2023-07-01

**Supplementary Figure 3.** Global prevalence of RBD convergent mutations (Fig. 5) across time. Data derived from Cov-spectrum[3]. Dashed lines correspond to the latest sampling data of a chronic-like clade where the respective mutations were detected**.**

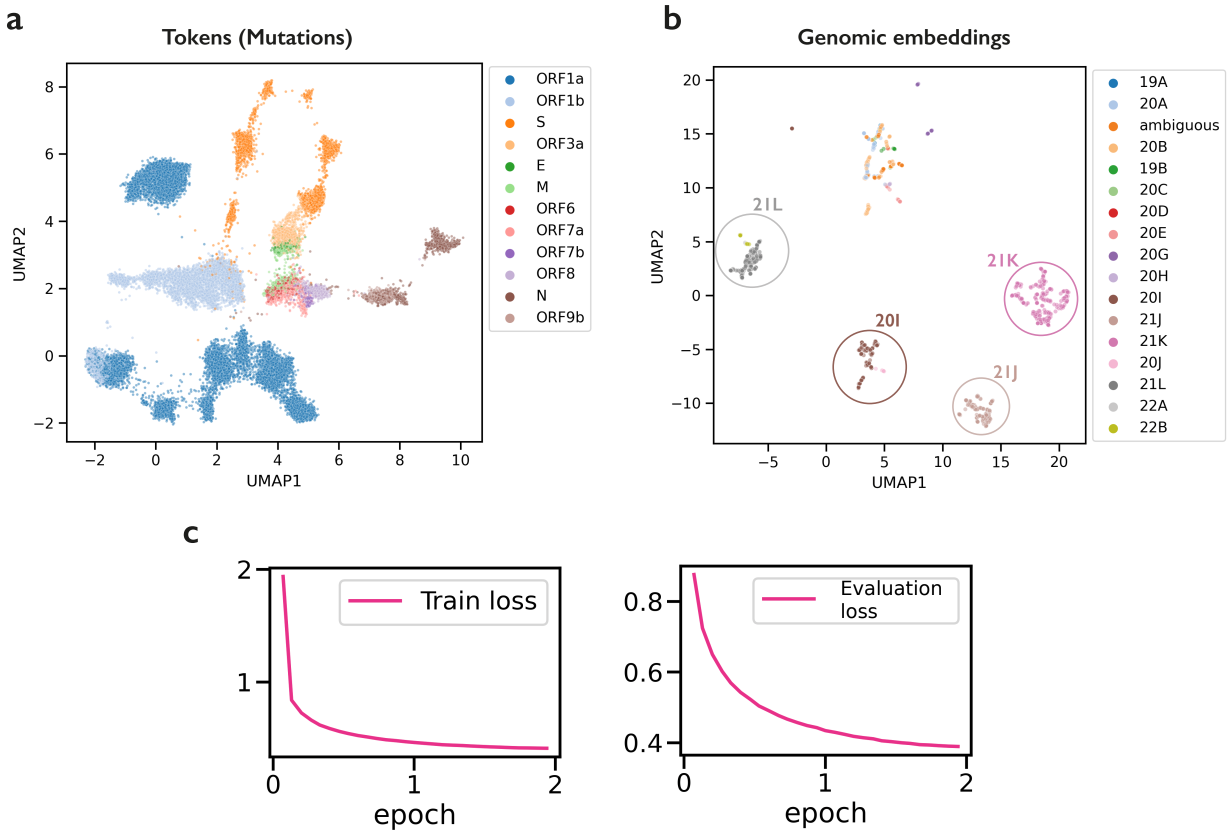


**Supplementary Figure 4. Language model embedding representation and training.** (a) Token (mutation) embedding projection into 2-dimensional space using UMAP of the trained BERT model. The mutations are color-coded by their genes. This analysis does not take into account the full genome sequence. (b) Genomic embeddings of all sequences of the 271 chronic-like clade (1337 sequences). The genome embedding was calculated by averaging the embeddings of each token of each sequence. This figure underscores the importance of the genomic embeddings for further biological predictions. (c) Train and Evaluation loss across 2 training epochs.

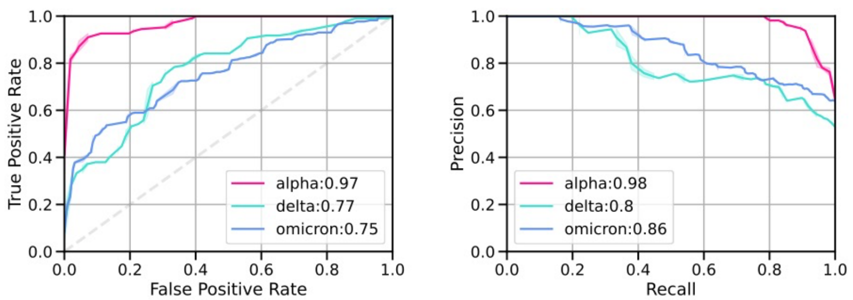


**Supplementary Figure 5. Classification performance assessment.** ROC and precision recall curves separated by the variant fold in the time and variant cross validation. The values described in the legend are the area under the curve for both ROC and precision recall graphs.
